# NANOTAXI: A Shiny-Based GUI for Real-Time Classification and Analysis of 16S rRNA Nanopore Reads

**DOI:** 10.64898/2026.05.17.725747

**Authors:** Nirmal Singh Mahar, Khushi Chouhan, Ishaan Gupta

## Abstract

Real-time taxonomic classification of nanopore amplicon sequencing data enables rapid insights into microbial communities, with applications in clinical diagnostics, environmental monitoring, and outbreak surveillance. However, bridging the gap between long-read data and interpretable results often requires specialised bioinformatics expertise. There remains a need for integrated, user-friendly software that combines live data acquisition with downstream microbiome analysis.

Here we present NANOTAXI, a fully automated Shiny-based GUI for the classification of barcoded 16S rRNA gene sequences generated by Oxford Nanopore sequencing. The platform supports four taxonomic classifiers, integrated with five reference databases, enabling flexible selection of classification strategies based on user requirements and available computational resources. In addition to real-time monitoring, NANOTAXI performs cohort-level analyses, including alpha and beta diversity, ordination, differential abundance testing, and functional inference using PICRUSt2.

Validation using barcoded synthetic communities comprising pooled genomic DNA from clinically relevant bacterial species and the ZymoBIOMICS mock community demonstrated that NANOTAXI generated biologically coherent taxonomic and functional profiles. Benchmarking revealed clear trade-offs between computational performance and taxonomic specificity. Emu provided the lowest observed species-level false-positive rate, whereas Kraken2 offered the fastest classification and enabled continuous near-real-time monitoring across all tested databases.

NANOTAXI is open source and freely available at https://github.com/Nirmal2310/NANOTAXI under the GPL version 3 license.

## 1. Introduction

High-throughput sequencing, often known as short-read sequencing, has revolutionised the study of microbial communities by providing high-quality data at a lower cost^1^. Another advantage of high-throughput sequencing, specifically in the context of the microbiome, is its ability to capture pathogenic bacteria that traditional culture-based techniques cannot^2^.

Compared to metagenomic sequencing, amplicon sequencing of the 16S rRNA gene has proven to be a reliable and efficient method for capturing microbial community composition^3–5^. Amplicon sequencing offers several advantages over metagenomic sequencing, including lower cost and greater sensitivity to host genome contamination^6^. The 16S rRNA gene contains both constant and variable regions: the constant regions are universally conserved across bacterial genomes, whereas the nine variable regions differ across bacterial genomes with varying degrees of variability^7^. Short-read sequencing typically targets and amplifies either a single variable region, such as V4 or V6, or three variable regions, such as V1-V3 or V3-V5, due to read length limitation^8^. However, targeting selective regions leads to uncertain taxonomic classifications for some bacterial genomes in complex microbial communities^9,10^.

Additionally, since data generation is not real-time, it can increase the overall data processing time, thereby delaying the mode of treatment^11^. Third-generation sequencing, or long-read sequencing, can overcome both these challenges by generating data in real time and sequencing the full-length 16S rRNA sequence, thereby improving species-level classification^12–14^. The ability of long-read sequencing, specifically Oxford Nanopore Sequencing (ONT), to generate data in real time has dramatically decreased the overall turnaround time to less than 6 hours, compared to the 48-72 hours required by traditional culture approaches, which is particularly relevant for clinical and field settings^15,16^.

The previously published tool NanoRTax^17^ demonstrated the feasibility of real-time taxonomic and diversity analysis for nanopore 16S rRNA amplicon data. However, the classifier, database and ONT software ecosystems have evolved substantially since its release, and existing workflows remain limited in terms of GUI-based accessibility, classifier/database flexibility, cohort-level statistical analysis and functional inference.

To address these challenges, we present NANOTAXI, an R Shiny-based GUI platform for analysing full-length barcoded 16S Nanopore sequencing data in real time. The platform allows users to analyse the barcoded data in both real-time and post-run (offline) mode. It integrates multiple widely used and recently developed classifiers and curated 16S reference databases within a single GUI-based workflow and performs both barcode-level and cohort-level microbiome analysis using best practices.

## 2. Materials and Methods

### 2.1 Overview of NANOTAXI

NANOTAXI is implemented as an R Shiny^18^ GUI that provides a simple, one-click interface for analysing barcoded long-read amplicon sequencing data. It integrates multiple widely used and recently developed classifiers and curated 16S reference databases within a single GUI-based workflow (**Figure 1**).

**Figure 1:**
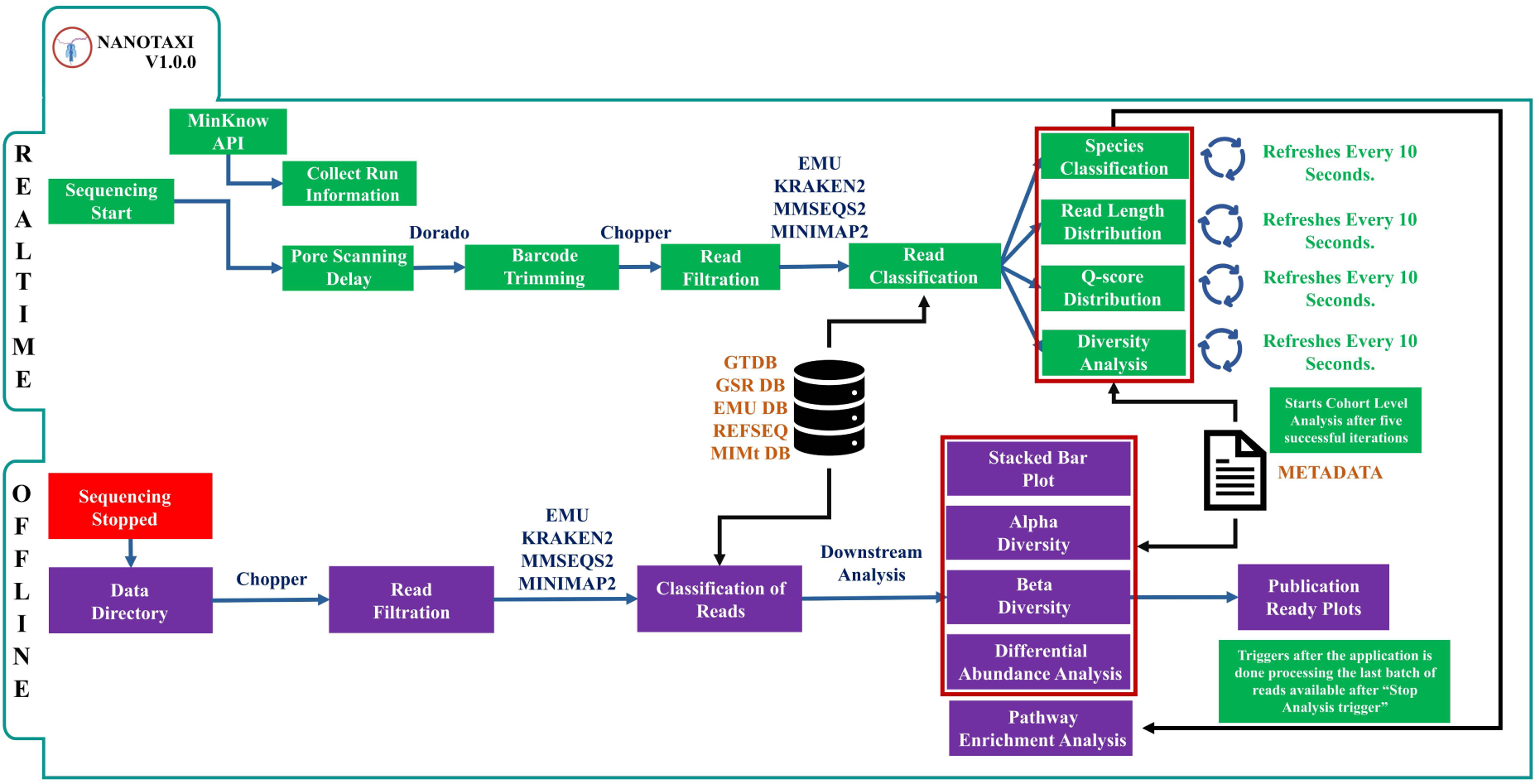
Overview of NANOTAXI. The application offers two separate routes for analysing sequencing data. In real-time mode, NANOTAXI connects to the sequencing device via the MinKNOW API to collect sequencing information for downstream analysis. Reads generated in real-time are pre-processed to retain full-length or near full-length rRNA sequences, which are then processed by the classification tool. Read counts per taxon are used to perform cohort-level analysis based on a user-provided metadata file. For post-run/offline mode, the user provides a directory containing barcoded reads. A similar classification approach is applied to these reads, and cohort-level analysis is performed on the resulting classifications.

Designed for ease of use, NANOTAXI enables researchers and clinicians without specialised bioinformatics training to process data in real time during a sequencing run or analyse existing data after the run. It allows users to export all generated visualisations as high-resolution PDF files suitable for reporting and publication. Thanks to its support for a wide range of classification tools and databases, NANOTAXI is not limited to clinical applications; it is also well-suited for environmental and forensic research.

### 2.2 Architectural design and core functionality

Using basecalled and demultiplexed reads, along with a comma-separated values (CSV) metadata file, NANOTAXI offers two distinct routes: one for real-time (online) data analysis and another for post-run (offline) analysis.

#### 2.2.1 Real-time mode

During real-time sequencing, NANOTAXI connects directly to the live run via the MinKNOW API (https://github.com/nanoporetech/minknow_api). This connection provides essential run metadata, including the output directory, number of active pores, sequencing status, and the kit used. This information allows the application to locate live data streams, monitor run progress, and apply kit-specific parameters for adapter and primer trimming.

Once pore scanning is complete, NANOTAXI begins classifying incoming reads. By default, it allocates 24 computational threads for parallel processing. These threads are distributed dynamically across active barcodes; for example, if only four barcodes contain new reads, each receives six threads. To ensure responsive performance, the system processes reads in batches of 500 reads per barcode per iteration. Users can adjust this “Batch Size”, though excessively large values may introduce lag depending on the pipeline and database in use (Supplementary Figure 1).

Between processing cycles, NANOTAXI pauses for 10 seconds (configurable via the “Update Interval” parameter) to prevent system overload. After five consecutive batches have been processed, the application automatically performs cohort-level analysis using the provided sample metadata. The cohort-level analysis auto-updates every 2 minutes to incorporate any new classified data.

When the user stops the analysis via the “Stop Analysis” button, NANOTAXI completes any ongoing batch before finalising the run. It then executes a functional profiling module powered by PICRUSt2^19^, which predicts functional gene abundances from the amplicon data. These results are subsequently available for differential abundance analysis and downstream interpretation.

#### 2.2.2 Offline/post-run mode

The offline analysis mode processes previously generated sequencing data. Analysis requires two primary inputs: a) a directory containing demultiplexed FASTQ files, structured with one sub-directory per barcode, matching the live-run output format of MinKNOW, and b) a corresponding metadata file, matching the format used in the real-time mode. At present, the application utilises only the first two columns: the first must specify the barcode name, and the second must define the sample’s categorical group for comparative analysis. Data processing begins with taxonomic classification, performed using a user-selected classification method and a reference database. Following classification, the resulting count matrix serves as the input for downstream cohort-level analyses.

Concurrently, the application executes a functional profiling module using PICRUSt2, similar to real-time mode. To ensure a responsive interface, all computationally intensive tasks, including the functional profiling performed by PICRUSt2, differential abundance analysis and real-time processing, are executed in separate R sessions. This prevents the main application from freezing during heavy computational tasks.

#### 2.2.3 Core analysis pipeline

##### 2.2.3.1 Data Preprocessing

Adapter and primer sequences from the demultiplexed reads are trimmed using the Dorado basecaller (https://github.com/nanoporetech/dorado). The trimming step is explicitly performed to reduce unclassified reads during the run, as trimming adapter sequences during demultiplexing has been reported to increase the rate of unclassified reads. The clean reads are then filtered using Chopper version 0.11.0^20^ to keep the high-quality reads. By default, reads with a Quality Score greater than or equal to 10 and length between 1400 and 1800 base pairs (bp) are retained to keep high-quality near full-length 16S rRNA gene sequences. However, users can modify these values at the start of the run via the GUI. The quality score distribution and read length distribution are determined using Bioawk version 20110810 (https://github.com/lh3/bioawk) and the readlength.sh module of BBMap version 37.62^21^, respectively.

##### 2.2.3.2 Taxonomic Classification

The high-quality, full-length or near full-length amplicon reads from the previous step are analysed for taxonomic classification. The current version of NANOTAXI supports 20 unique combinations of classification methods and databases (**Table 1**). Supported methods include Minimap2 version 2.30^22^, Kraken2 version 2.17.1^23^, Emu version 3.4.5^24^ and MMseqs2 version 18.8cc5c^25^, which are packaged with five reference databases named NCBI RefSeq 16S^26^, GSR^27^, GTDB^28^, Emu proprietary database^24^ and MIMt^29^, to ensure wide applicability beyond clinical samples. Multiple classification strategies are incorporated to accommodate differences in computational requirements, taxonomic resolution and analysis objectives. Therefore, the application allows users to select the appropriate tool-database combination based on their specific requirements, whether for rapid monitoring or high-resolution, computationally intensive analysis.

**Table 1:** Read classification methods and databases utilised by NANOTAXI for both real-time and offline modes. At present, NANOTAXI supports all combinations of classification methods and databases in both modes.

| Classifier | Mode | RefSeq 16S | GSR-DB | MIMt | EMUDB | GTDB |
| --- | --- | --- | --- | --- | --- | --- |
| Kraken2 | Both | ✓ | ✓ | ✓ | ✓ | ✓ |
| Minimap2 | Both | ✓ | ✓ | ✓ | ✓ | ✓ |
| Emu | Both | ✓ | ✓ | ✓ | ✓ | ✓ |
| MMseqs2 | Both | ✓ | ✓ | ✓ | ✓ | ✓ |

The initial classification output is filtered according to user-defined parameters set at the start of the analysis. These include minimum percentage identity, minimum percentage coverage, and a confidence score (for Kraken2 results). Following the filtration step, the refined output is processed with Taxonkit version 0.17.0^30^ to retrieve a complete, standardised taxonomic lineage for each identified taxon using its native database. This full taxonomy is subsequently used for all downstream analysis and visualisation in the application.

##### 2.2.3.3 Data curation

After taxonomic classification, NANOTAXI compiles the relevant output files, including per-barcode data on read length distribution, read quality scores, and classification results. The length and quality data are immediately plotted as a bar chart, providing users with real-time, barcode-specific feedback on sequencing performance, unlike MinKNOW, which aggregates these metrics across the entire run. This feature enables informed decisions about whether to continue or terminate a sequencing run based on the quality of individual samples.

The classification data is used to generate several analytical plots. A rarefaction curve displays the cumulative number of unique taxa observed over time, helping assess whether sample diversity has been fully captured, as indicated by a plateau. By default, only species with at least 10 supporting reads are included, though this threshold can be adjusted via the “Absolute Counts Cutoff” parameter.

Additionally, the system calculates two key diversity indices, the Shannon index, which emphasises species richness and rare taxa and the Simpson index, which is sensitive to dominant species. Both these metrics are utilised to track the community evenness and stability as the run progresses. Together, the rarefaction and diversity plots inform users when sufficient data has been collected, guiding decisions to stop sequencing efficiently.

Finally, classification results from all barcodes are merged into a unified taxa count table, which serves as the basis for all downstream cohort-level analysis.

#### 2.2.4 Cohort-level analysis

For cohort analysis, we employed best practices for microbiome data analysis outlined by Gloor et al.^31^, including compositional data transformations and appropriate statistical methods.

##### 2.2.4.1 Compositional visualisation

Compositional visualisation analysis begins with the Total-Sum Scaling (TSS) normalisation of the unified taxa counts, generating relative abundance profiles for each barcode. These profiles are visualised in an interactive stacked bar plot, where the x-axis represents individual barcodes and the y-axis shows relative abundance (%). This plot serves as the foundational step in cohort-level analysis, enabling users to rapidly assess sample similarity, identify dominant taxa, and detect major compositional shifts with metadata groupings.

By default, the plot displays the top five most abundant taxa across all barcodes, with all remaining taxa grouped into an “Other” category. Users can dynamically customise this view by adjusting the “Top N” slider to display up to 25 top taxa. Furthermore, the plot can be generated for any taxonomic level, from phylum to species, selected in the “Taxon Level” option. To focus on a specific organism of interest, users may enter a taxon name in the “Select Taxon Name” field, after which the plot will display only the taxon’s relative abundance across barcodes.

Alongside the visualisation, NANOTAXI provides the underlying unified taxa count table for review under the “Analysis Results” tab within the “INPUT” tab. This table can be downloaded as a tab-separated value (.tsv) file, ensuring compatibility with downstream analysis in other bioinformatics platforms.

##### 2.2.4.2 Alpha diversity analysis

Alpha diversity measures the complexity of a microbial community within a single sample. To enable accurate comparisons across samples with varying sequencing depths, NANOTAXI performs rarefaction. This process randomly subsamples each sample to an equal read count, thereby controlling for biases introduced by differences in library size.

The rarefaction threshold is determined using Good’s coverage index (Equation 1), a metric that estimates sampling completeness. The application selects the minimum library size that achieves a Good’s coverage of 95% as the subsampling cutoff.

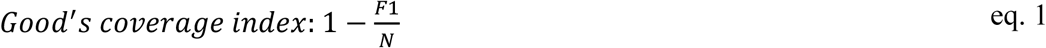

F1 = number of singleton operational taxonomic units (OTU)

N = Sum of counts for all OTUs.

The OTU count matrix is rarefied to this cutoff for 100 iterations. Alpha diversity metrics (Shannon and Simpson indices)^32^ are calculated at each iteration, and their average value is computed per sample. Samples are grouped by metadata, and the Wilcoxon rank-sum test is used to evaluate differences in alpha diversity across groups. The resulting p-values are corrected for multiple testing using the Benjamini-Hochberg (BH) method.

The final output is presented as a box plot showing the distribution of each alpha-diversity metric across groups. Groups with a statistically significant difference (adjusted *p*-value < 0.05) are marked with an asterisk (*).

##### 2.2.4.3 Beta diversity analysis

Beta diversity analysis assesses differences in microbial community composition between samples. Two parallel pipelines process the unified taxa count table to account for the compositional nature of microbiome data.

The count data is first normalised using Total-Sum Scaling (TSS) to generate relative abundance profiles. To reduce noise, taxa are filtered by a minimum prevalence threshold (default: 10% of barcodes) and a minimum mean relative abundance threshold (default: 0.1%); both thresholds are user-adjustable. The filtered data is renormalised via TSS. Bray-Curtis’s dissimilarity is calculated on this matrix to quantify compositional differences between samples. Non-metric Multidimensional Scaling (NMDS) is then applied to the resulting distance matrix, preserving the rank order of barcode dissimilarities. The NMDS ordination is visualised as a scatter plot, with barcodes positioned along the first two NMDS axes and coloured by group information from the metadata.

To address the constant-sum constraint inherent in relative abundance data, a centred log-ratio (CLR) transformation is applied. A pseudo-count of 1e^-10^ is added to enable log-transformation of zero values. The CLR-transformed data is further used for two additional ordination methods: Principal Coordinate Analysis (PCoA) based on Euclidean distance (equivalent to the Aitchison distance for compositional data), with results displayed as a scatter plot of the first two principal coordinates, annotated with their explained variance. Principal Component Analysis (PCA), which visualises multivariate compositional differences. The resulting biplot overlays the top taxa; the default is 5, but it can be adjusted to 25 via the “Biplot Top Taxa” slider, highlighting the taxa driving group separation.

Group differences in community composition are statistically evaluated using Permutational Multivariate Analysis of Variance (PERMANOVA) with Euclidean distance on the CLR-transformed data and 9,999 permutations. Results are presented in a table showing pairwise group comparisons, *R*^2^, *p*-value, and Benjamini-Hochberg-adjusted *p*-value (*p*_adj_), available for download as a tsv file.

Finally, a heatmap is generated from z-score-standardised CLR-transformed values using the ComplexHeatmap R package^33^. Rows (taxa) and columns (barcodes) are clustered using hierarchical clustering with Euclidean distance and complete linkage. Group annotation for each barcode is derived from metadata, and taxa prevalence is displayed as a right-side bar plot annotation.

Overall, for beta diversity, NANOTAXI employs Bray-Curtis dissimilarity on TSS-normalised data and the Aitchison distance (via the CLR transformation) to address compositional data. Outputs include an NMDS scatter plot, PCoA scatter plot, PCA biplot, a PERMANOVA results table, and an annotated heatmap. Together, these complementary approaches provide a robust and comprehensive assessment of beta diversity within the NANOTAXI platform.

##### 2.2.4.4 Taxa Differential abundance analysis

To identify taxa with significantly different abundances between predefined sample groups, NANOTAXI performs differential abundance analysis using the Analysis of Compositions of Microbiome with Bias Correction 2 (ANCOM-BC2) method^34^, which is designed for compositional, sparse, and high-dimensional microbiome count data. The analysis is applied to the unified taxa count table, where rows represent taxa and columns represent barcodes.

Before analysis, the counts table is filtered using a prevalence cutoff (default: 10% of the samples) and a library-size cutoff (default: minimum of 10 reads per barcode) to remove rare taxa and low-depth samples. Both thresholds are user-adjustable via the “Prevalence Cutoff (%)” and “Absolute Counts Cutoff” parameters in the application’s COHORT tab.

ANCOM-BC2 is executed on the filtered data with a fixed effect of the form Abundance∼ Group, without random effects. Resulting *p*-values are corrected for multiple testing using the Benjamini-Hochberg (BH) procedure, and statistical significance is assessed at alpha = 0.05.

For all-vs-all comparisons, a pairwise directional test is applied to each pairwise comparison. The resulting table is used to identify differentially abundant taxa for each comparison.

Results for a selected comparison are visualised in a volcano plot, where the x-axis shows log_2_ fold change and the y-axis shows -log_10_(adjusted *p*-value). Taxa are classified as enriched (log_2_ fold-change ≥ 1 and adjusted *p*-value ≤ 0.05) or depleted (log_2_ fold-change ≤ −1 and adjusted *p*-value ≤ 0.05), and their labels are displayed on the plot. The comparison group can be selected via a dropdown menu in the DAA tab. The full pairwise directional test table is available for download as a TSV file for use in downstream analyses.

##### 2.2.4.5 Functional Inference

To infer the metabolic potential of observed microbial communities, NANOTAXI includes a functional prediction pipeline based on PICRUSt2 (Phylogenetic Investigation of Communities by Reconstruction of Unobserved States 2)^19^. This tool translates 16S rRNA gene taxonomic profiles into predicted abundances of gene families and metabolic pathways, enabling functional interpretation of microbiome data.

PICRUSt2 requires a taxon count table and a FASTA file containing representative sequences for each species. For species associated with multiple reference sequences, the medoid sequence is selected as the taxon representative. The medoid is determined by computing a pairwise distance matrix using the normalised Levenshtein distance, then selecting the biological sequence with the smallest cumulative distance to all other sequences within the same taxonomic group.

PICRUSt2 places the representative sequences into a reference phylogenetic tree using EPA-ng^35^. It then predicts gene families per taxon using ancestral-state reconstruction implemented in the Castor R package^36^. Predicted gene families are summed per sample to generate abundance tables for KEGG Orthologs (KOs) and Enzyme Commission (EC) numbers. Subsequently, the HUMAnN2 pipeline^37^ is used internally to map KO abundances to MetaCyc metabolic pathways via MinPath^38^, producing a pathway-level abundance table.

The resulting KO and MetaCyc abundance tables are normalised using Total Sum Scaling (TSS) and transformed using the Centred Log-Ratio (CLR). Principal Component Analysis (PCA) is performed on each transformed table, and the results are displayed as a scatter plot of the first two principal components, with the explained variance annotated. Users can select the functional category (KO or MetaCyc) using the “Functional Pathway Group” dropdown menu.

Differential abundance analysis is performed on all functional tables using ANCOM-BC2. Differentially abundant KEGG orthologs are further ranked by adjusted p-value (padj) and subjected to Gene Set Enrichment Analysis (GSEA) using clusterProfiler^39^. The top enriched pathways (10 by default, adjustable up to 20 per comparison) are displayed as a dot plot. For MetaCyc pathways, differentially abundant results are visualised as a bidirectional bar plot showing log₂ fold changes. All differential abundance results can be downloaded as a TSV file for further investigation.

#### 2.2.5 Implementation and availability

NANOTAXI is implemented as an R Shiny application that leverages R’s extensive ecosystem for bioinformatics, statistical analysis, and interactive visualisation. The software follows a modular architecture, with dedicated modules for real-time data processing, taxonomic classification, and each cohort-level analytical component (e.g., diversity, differential abundance, functional profiling).

Software dependencies, including all required R packages and external bioinformatics tools, are managed automatically via Conda and Pip. Upon first launch, the application installs and configures these dependencies, downloads the relevant databases, and ensures a streamlined setup for users without specialised computational expertise. A complete list of integrated tools and databases is provided in Supplementary Table 1.

NANOTAXI is released as open-source software under the GNU General Public License v3.0. The full source code, along with comprehensive documentation, installation guides, and example datasets, is publicly available on GitHub at: https://github.com/Nirmal2310/NANOTAXI

#### 2.2.6 Synthetic community sequencing

Reference strains obtained from the American Type Culture Collection (ATCC) were subcultured on blood agar plates and incubated at 37°C until isolated colonies were observed. Single colonies were inoculated into Luria-Bertani (LB) broth and incubated overnight to an optical density (OD600) of 0.8–1.2. Bacterial cells were harvested by centrifugation, and genomic DNA (gDNA) was extracted using the QIAamp DNA Mini Kit (Qiagen; cat. no. 51304) according to the manufacturer’s instructions. The purity and concentration of the extracted gDNA were assessed using a NanoDrop spectrophotometer.

To simulate mixed microbial populations with defined abundance profiles, four distinct mock communities (designated A-D) were constructed. Extracted gDNA from three species (*Escherichia coli, Pseudomonas aeruginosa, and Staphylococcus aureus*) and the ZymoBIOMICS community was mixed in equimolar mixtures and prepared in biological triplicates (barcode 01-03). For the remaining samples, each community was designed to be dominated by a single species (80% of total DNA), with the remaining two species serving as background populations (10% each) (**Table 2**). Finally, three biological replicates of the ZymoBIOMICS mock community were also included in the sequencing run (barcode 13-15).

**Table 2:** Detailed information regarding the composition of the synthetic microbiome communities created for testing the real-time functionality of NANOTAXI.

| Community | Composition | Barcodes |
| --- | --- | --- |
| **A** | *E. coli* (25%), *P. aeruginosa* (25%), *S. aureus* (25%),<br>ZymoBIOMICS (25%) | 01-03 |
| **B** | *E. coli* (80%), *P. aeruginosa* (10%), *S. aureus* (10%) | 04-06 |
| **C** | *E. coli* (10%), *P. aeruginosa* (80%), *S. aureus* (10%) | 07-09 |
| **D** | *E. coli* (10%), *P. aeruginosa* (10%), *S. aureus* (80%) | 10-12 |
| **ZymoBIOMICS** | Commercial mock community | 13-15 |

Sequencing libraries were prepared using the 16S Barcoding Kit 24 V14 (SQK-16S114.24, Oxford Nanopore Technologies) following the manufacturer’s standard protocol. Briefly, the 16S rRNA gene was amplified and barcoded for each triplicate sample, and the products were purified using magnetic beads. The purified barcoded libraries were quantified and pooled in equimolar ratios. The final library pool was loaded onto a SpotON Flow Cell (R10.4.1) and sequenced using a MinION Mk1D device.

## 3 Results

### 3.1 Real-time analysis

To validate NANOTAXI’s real-time analysis capability, we sequenced a series of barcoded synthetic communities composed of pooled genomic DNA from clinically relevant bacterial species, as well as the ZymoBIOMICS mock community. Each experimental community featured a distinct dominant species to simulate an infection, while a control community contained all species at approximately equimolar proportions. Three technical replicates per community were included in the final sequencing pool to mitigate technical noise and distinguish biological signals from artefacts.

NANOTAXI successfully established a live connection to the sequencing instrument via the MinKNOW API. The interactive dashboard updated dynamically, displaying critical run metrics including sequencing status, active pore count, processed reads, mean read length, and, for each barcode, the number of classified reads and unique taxa identified (**Figure 2**). This consolidated display provided all essential information within a single interface, eliminating the need for users to switch between NANOTAXI and the MinKNOW software.

**Figure 2:**
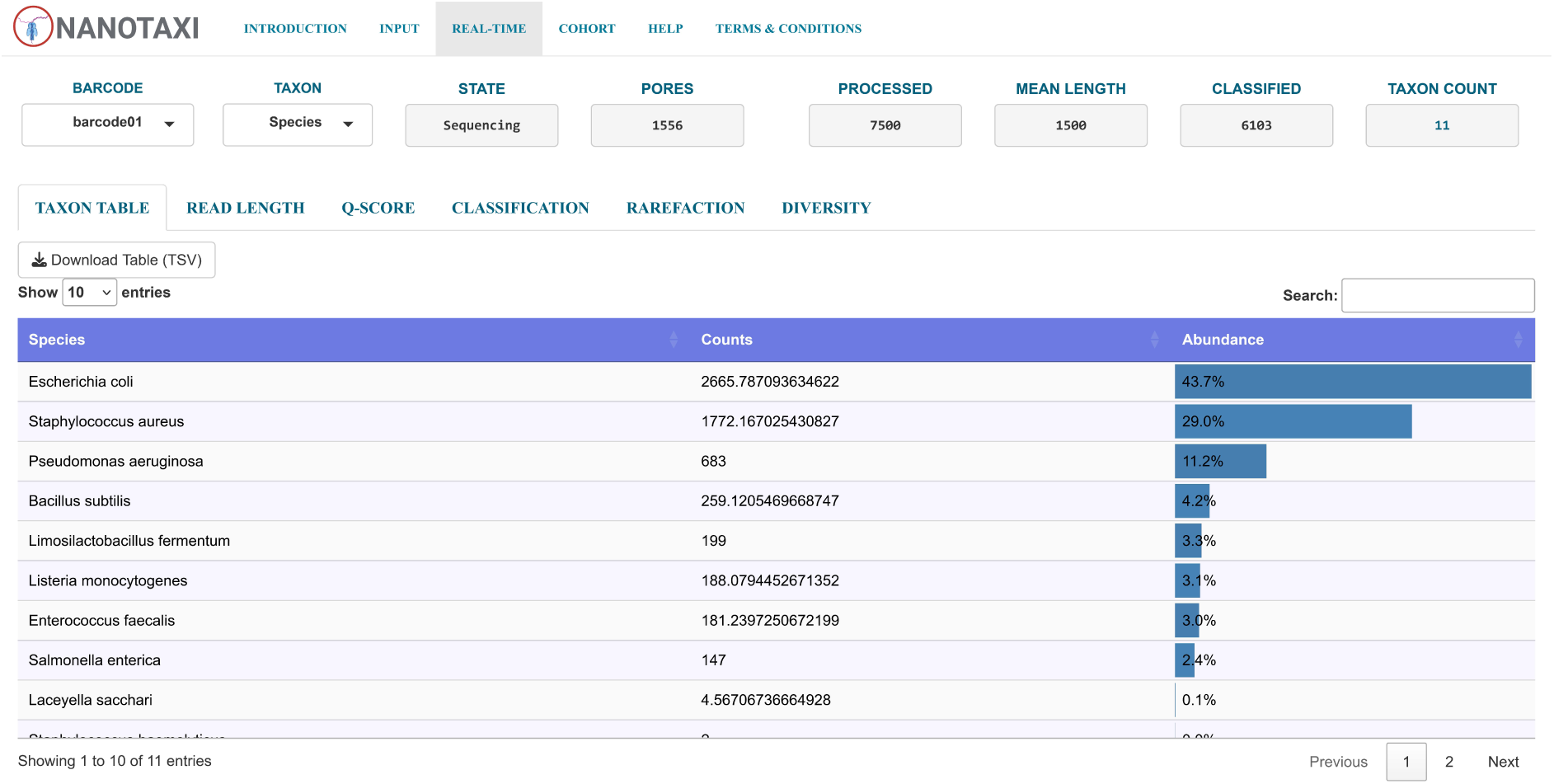
The real-time analysis interface of the NANOTAXI application. The application displays information about the live run, allowing users to avoid switching between the MinKNOW application and NANOTAXI. The interface offers multiple control widgets for real-time adjustments to the plots generated during the run. The current screenshot displays the taxon abundance table for barcode 01, which users can download as a TSV file. Users can display the generated plots for each available barcode and adjust the taxon level to kingdom level.

Sequencing quality was monitored in real time through per-barcode distributions of read length and quality score (Q-score). Reads within the target amplicon range (1,400–1,800 bp, representing full-length or near-full-length 16S rRNA gene sequences) and with a Q-score ≥ 10 were designated high-quality (cyan azure) and retained for analysis; all others (charm pink) were filtered out (**Figure 3A, B**). Throughout the run, the majority of reads met the high-quality threshold.

**Figure 3:**
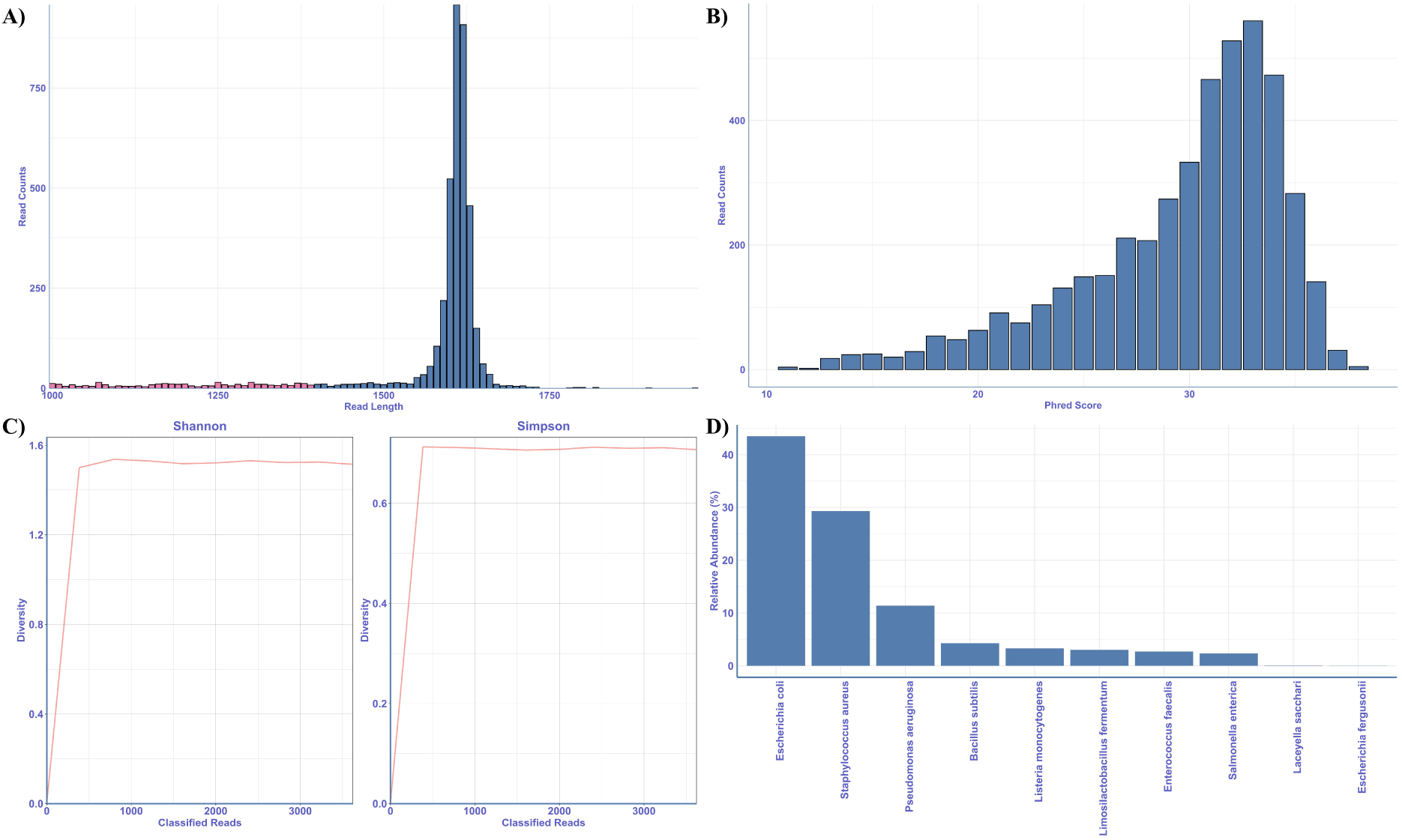
Real-Time Analysis and Monitoring in NANOTAXI. **A)** Bar plot showing the distribution of read lengths for each barcode. Reads within the target amplicon range (1,400–1,800 bp, representing full-length or near-full-length 16S rRNA gene sequences) are shown in cyan azure. Reads outside this range are coloured in charm pink. B) Bar plot displaying the distribution of quality scores (Q-scores). Reads with Q ≥ 10 (high quality) are shown in cyan azure; reads with Q < 10 (higher error rate) are shown in charm pink. C) Line plot tracking Shannon and Simpson diversity indices calculated from classified reads as the run progresses. The Shannon index emphasises species richness and rare taxa, while the Simpson index is more sensitive to the dominance of common species. D) Bar plot of the top 10 most abundant species (or other selected rank) for a selected barcode, displayed as either read counts or relative abundance. Users can adjust the display to show up to the top 25 taxa at any taxonomic level (kingdom to species). All plots are available for each barcode and can be downloaded as PDFs for reporting.

These high-quality reads were classified taxonomically using Emu with its native database on a system equipped with an AMD EPYC 7543 32-core processor and 314 GB of RAM. Classification results were displayed as an interactive bar plot. For example, after 1 hour of sequencing, the platform had processed ∼10,000 reads for barcode 01 (Equimolar community). The composition was dominated by *Escherichia coli* (43.5%), followed by *Staphylococcus aureus* (29.3%), *Pseudomonas aeruginosa* (11.2%), *Bacillus subtilis* (4.37%), *Listeria monocytogenes* (3.17%), *Lactobacillus fermentum* (3.12%), *Enterococcus faecalis*(2.85%) and *Salmonella enterica* (2.36%) (**Figure 3D**). NANOTAXI detected all expected members in every community. However, observed abundances deviated from the expected theoretical proportions.

The platform also calculated alpha diversity (Shannon and Simpson indices) from the classified reads in real time. This feature allows users to determine the point of diminishing returns in sequencing depth, as indicated by a plateau in the diversity curve. For barcode 01, the curve plateaued after approximately 1,000 classified reads, suggesting that the full diversity of the community had been captured (**Figure 3C**). sequencing was continued to ensure sufficient coverage for all remaining barcodes.

### 3.2 Cohort-level analysis

NANOTAXI automatically triggered cohort-level analysis after five successful classification iterations; however, the results presented here are based on the complete dataset generated at the end of the sequencing run.

Using the metadata file provided at the outset, the platform performed all major group-level comparisons. It includes interactive control widgets that allow users to customise output visualisations and filter input data for each analysis (**Figure 4**). Adjustable parameters include prevalence, relative abundance, and library size cutoffs, among others. This customisable functionality, a feature not commonly available in comparable platforms, enables researchers to tailor the analysis to their specific experimental context and data characteristics.

**Figure 4:**
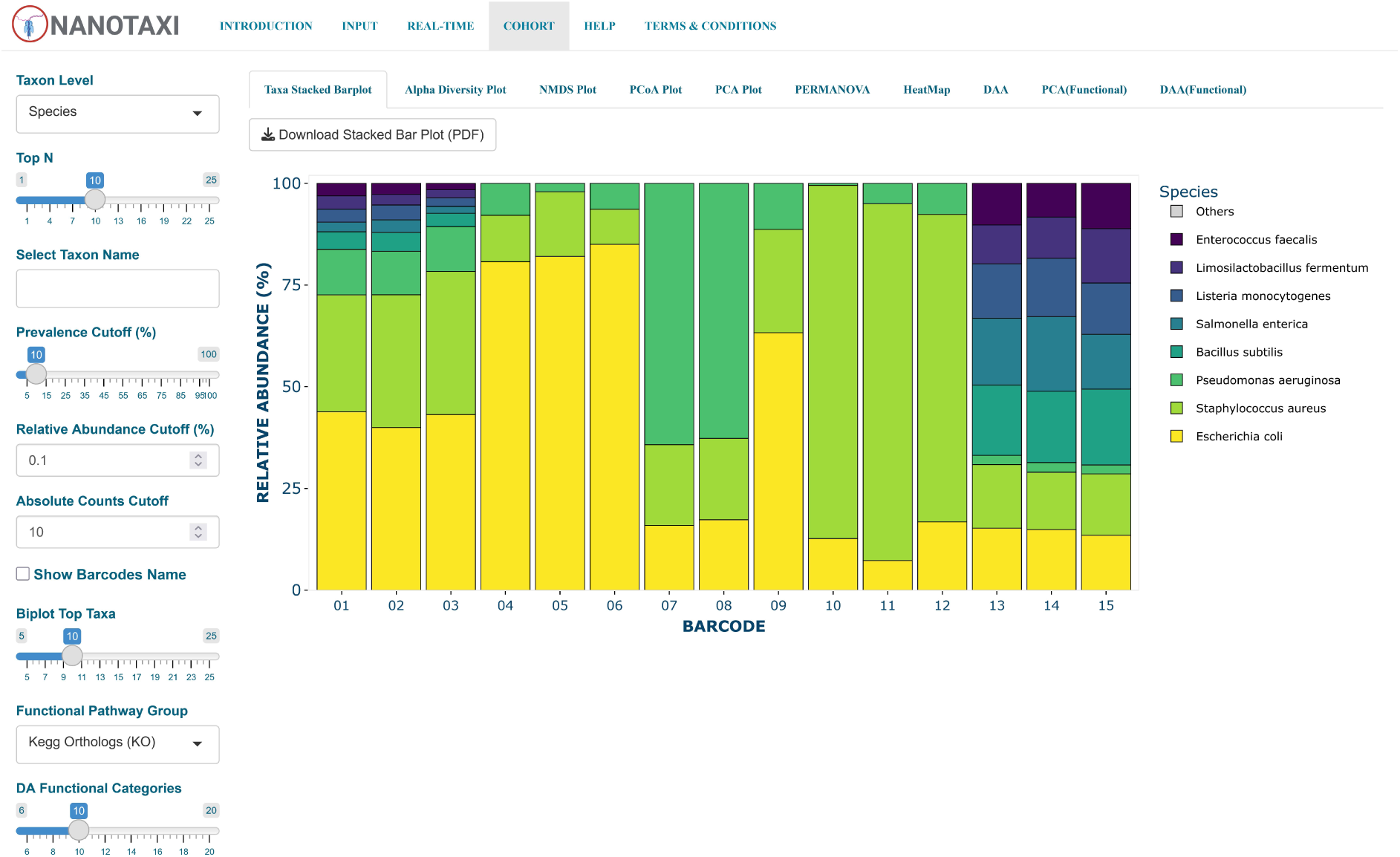
The cohort analysis interface of the NANOTAXI application. The left sidebar contains control widgets for selecting taxonomic levels and setting data-filtration parameters, such as prevalence, relative abundance, and absolute count cutoffs. The control widgets allow users to make real-time adjustments to the group-level comparative analysis of the amplicon data.

The community composition for each barcode was visualised using a stacked bar plot of the top ten species. The analysis confirmed the pattern observed during real-time classification: *Escherichia coli* and *Staphylococcus aureus* were consistently dominated over *Pseudomonas aeruginosa* across all barcodes categorised under equimolar group compared to its expected proportion (**Figure 5A**). In the equimolar group (barcodes 01–03), *E. coli* had a mean relative abundance of 42.1% against an expected 25%, *S. aureus* had a mean relative abundance of 32.2% and *P. aeruginosa* had a mean relative abundance of 11.1%. Additionally, barcode 09, which was categorised under the *Pseudomonas aeruginosa* dominant group, had *Escherichia coli* as the dominant species with 63.8% relative abundance. This unexpected observation might be due to cross-contamination during library preparation.

**Figure 5:**
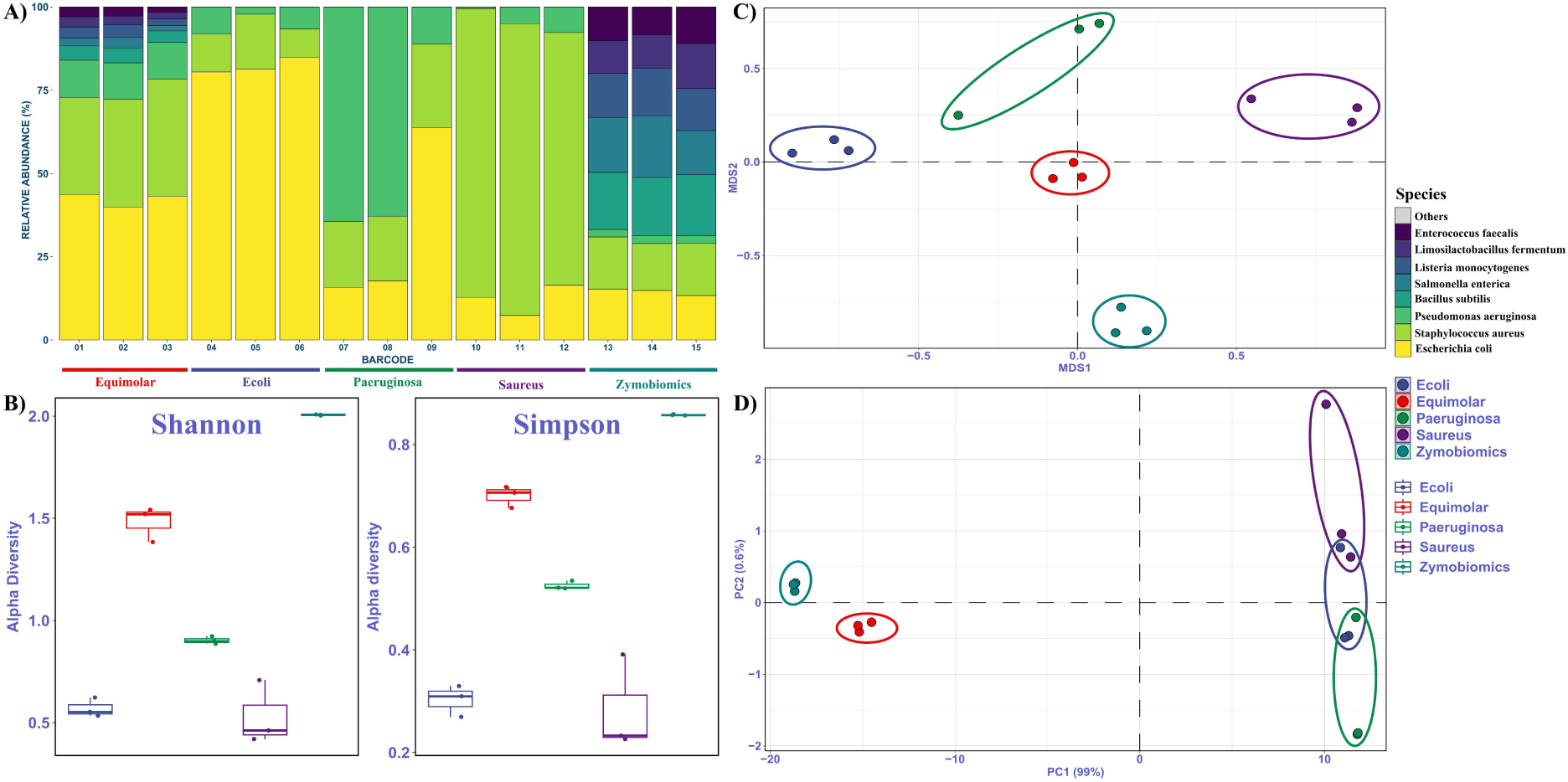
Cohort-level analytical outputs generated by NANOTAXI. Plots derived from the unified taxa count table and user-provided metadata. **A)** Stacked bar plot displaying the relative abundance of the top five most abundant species per barcode; remaining taxa are grouped as “Others”. Users can adjust the display to show the top 25 taxa at any taxonomic rank (phylum to species) and can subset the view to track a specific taxon across samples. **B)** Box plots of Shannon and Simpson indices within-sample diversity (richness and evenness) across experimental groups. Statistical significance is assessed using the Wilcoxon rank-sum test, with significant pairwise differences (adjusted p-value < 0.05) indicated by asterisks and connecting brackets. **C)** Non-metric multidimensional scaling (NMDS) ordination based on Bray-Curtis dissimilarity of total-sum-scaled (TSS) abundance data, visualising sample clustering and inter-group distances. **D)** Principal Coordinate Analysis (PCoA) ordination plot of the first two coordinates, using Aitchison distance (Euclidean distance on CLR-transformed data). The percentage variance explained is displayed on each axis.

Within-sample diversity was assessed for each group using the Shannon and Simpson indices, and the results were displayed as a grouped box plot (**Figure 5B**), aligned with compositional expectations. The *Staphylococcus aureus*–dominant community exhibited the lowest alpha diversity, consistent with its high average relative abundance of 83.4%, closely followed by the *Escherichia coli*-dominant community with an average relative abundance of 82.2%. As expected ZymoBIOMICS synthetic community showed the highest diversity, followed by the equimolar group. However, the pairwise Wilcoxon rank-sum test indicated that none of the observed differences in alpha diversity across groups was statistically significant.

Beta diversity analysis was conducted to evaluate compositional differences between the synthetic bacterial communities. While ordination plots indicated moderate separation among groups, these differences were not statistically significant according to PERMANOVA.

Non-metric multidimensional scaling (NMDS) based on Bray-Curtis dissimilarity revealed distinct clustering, with barcode 09 from the *P. aeruginosa* dominant group positioned closest to the *Escherichia coli* dominant group, as expected from its compositional profile (**Figure 5C**).

Principal Coordinate Analysis (PCoA) using Aitchison distance reinforced the separation of the ZymoBIOMICS and Equimolar communities, which together explained 99.6% of the variance along the first two coordinates (**Figure 5D**). In contrast to the NMDS plot, the remaining communities did not form distinct clusters in the PCoA ordination with the first two principal coordinates. Therefore, we further demonstrated community clustering in the second and third principal coordinates (Supplementary Figure 2). The resulting ordination plot clearly segregated the communities, with barcode 09 positioned closest to the *E. coli*-dominant group, similar to the NMDS ordination plot.

Additionally, a Principal Component Analysis (PCA) biplot based on the CLR-normalised abundance data further identified the taxa driving the clustering of the groups (Supplementary Figure 3). Barcodes from the ZymoBIOMICS group clustered closer to the origin, indicating a balanced composition. Species unique to the ZymoBIOMICS synthetic community were strongly associated with the separation of the ZymoBIOMICS group along the first principal component. The equimolar group, *E. coli*-dominant group, and *S. aureus*-dominant group clustered on the positive side of the first component, with *E. coli* and *S. aureus* as the primary drivers. The *P. aeruginosa* dominant group was separated on the negative side of the second component, with P. aeruginosa as the main taxon driving the clustering. Overall, the PCA confirmed that each group’s intended dominant species contributed to its separation.

Finally, a clustered heatmap of z-score-standardised CLR-transformed abundances provided a taxon-level overview of community structure (Supplementary Figure 4). Hierarchical clustering grouped samples largely by their expected dominant species, with barcode 09 clustering with the *E. coli* dominant group.

NANOTAXI includes taxon differential abundance analysis as part of its cohort-level analysis suite. While it performs this analysis in an all-vs-all manner, we have incorporated results specifically from the comparison between the in-house synthetic communities and the ZymoBIOMICS mock community.

The results validated the expected compositional patterns for the communities: in the *dominant groups Escherichia coli, Staphylococcus aureus, and Pseudomonas aeruginosa*, the intended dominant species was significantly enriched relative to the control (**Figure 6A, B, C**). In the equimolar community, all three introduced species were significantly enriched relative to the ZymoBIOMICS reference community (**Figure 6D**).

**Figure 6:**
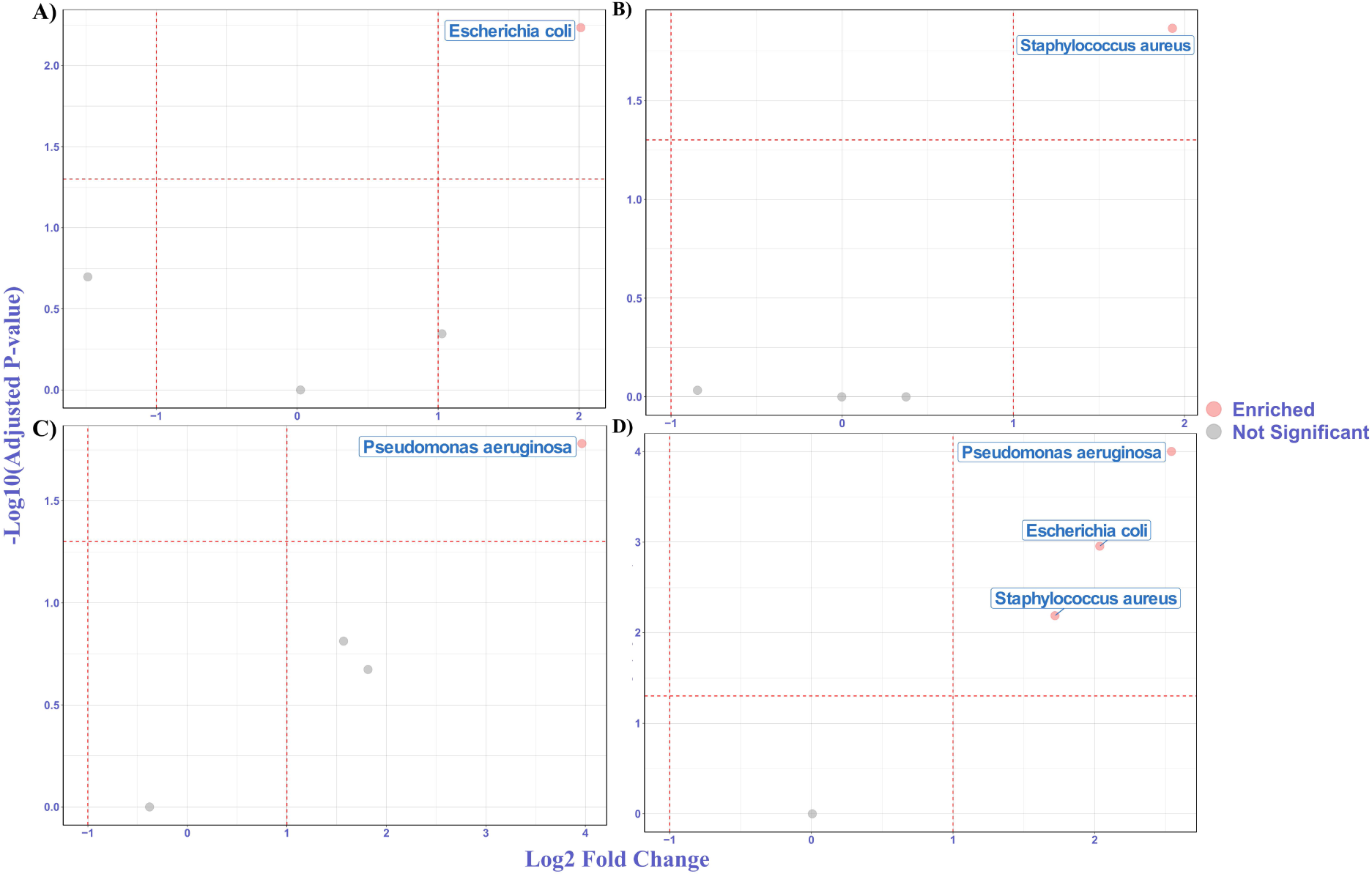
Volcano plots comparing each synthetic community against the ZymoBIOMICS reference community. A) *Escherichia coli* dominant community. B) *Staphylococcus aureus* dominant community. C) *Pseudomonas aeruginosa* dominant community. D) Equimolar community. In each panel, species are plotted by log2 fold change versus -log_10_(adjusted *p*-value). Significantly enriched or depleted species are labelled.

Following the final classification cycle, NANOTAXI executed its integrated functional inference module, PICRUSt2, on the synthetic community dataset.

Principal Component Analysis (PCA) of predicted KEGG Ortholog (KO) abundances revealed a clear separation of the groups along the second principal component (**Figure 7A**), with barcode 09 showing a pattern similar to that observed in the species abundance-based ordination. A similar separation pattern was observed in the PCA of the MetaCyc pathways abundance (**Figure 7C**). These results demonstrate the platform’s capacity to propagate taxonomic differences into functional feature space.

**Figure 7:**
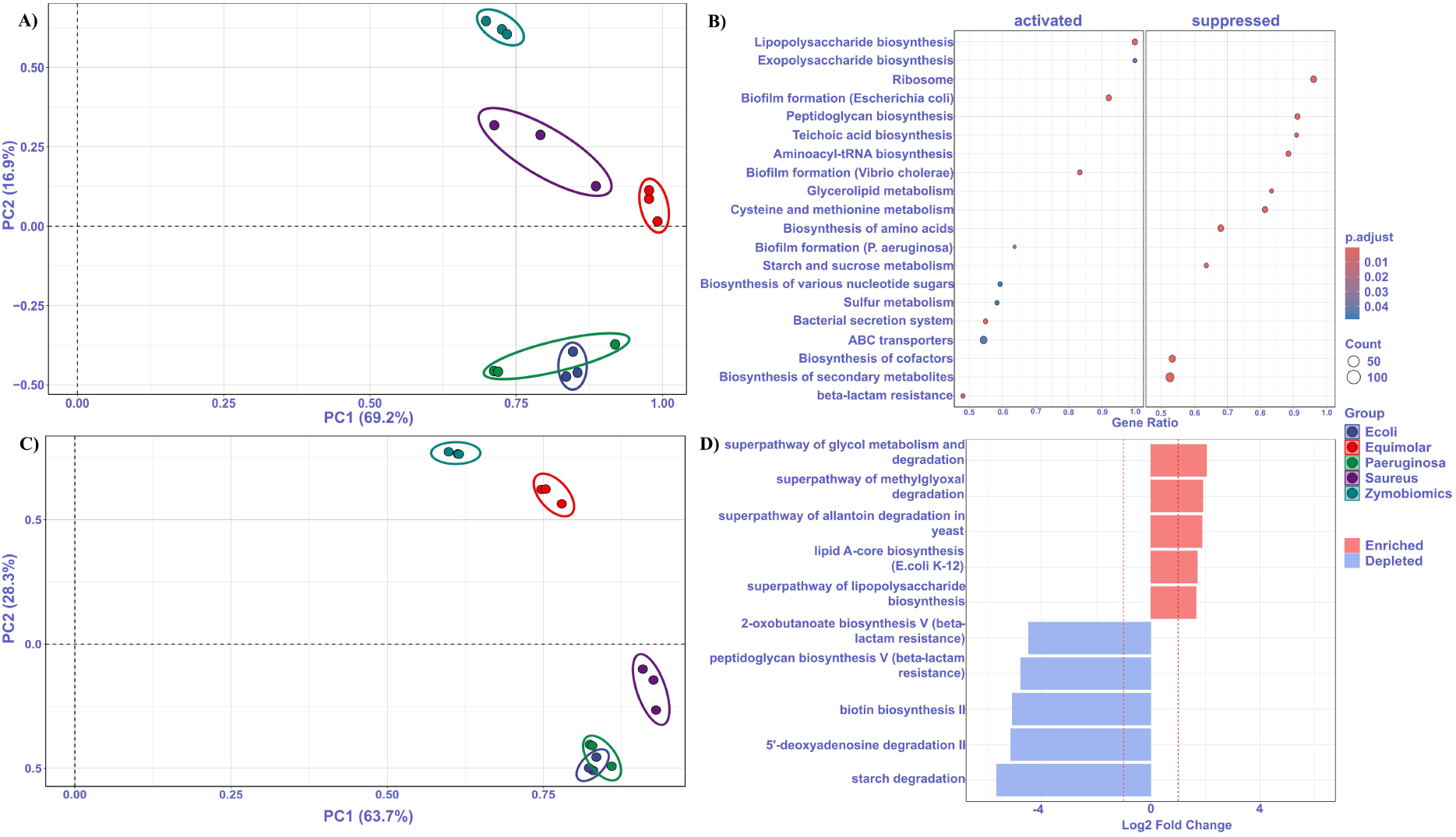
Functional inference and Enrichment analysis from PICRUSt2. **A)** PCA ordination of barcodes based on KEGG Ortholog (KO) abundance. The percentage of variance explained by each principal component (PC) is shown along the axes. **B)** Dot plot showing the enriched and depleted predicted pathways associated with the differentially abundant KOs in the *Escherichia coli* dominant group in comparison to the ZymoBIOMICS group. **C)** PCA ordination of barcodes based on MetaCyc pathways abundance. The percentage of variance explained by each principal component (PC) is shown along the axes. **D)** Bidirectional bar plot of differentially abundant MetaCyc pathways in the *Escherichia coli* dominant group with respect to the ZymoBIOMICS group. Bar length indicates log2 fold-change; colour denotes pathway category. Equivalent plots are generated for all comparisons.

Similar to the approach used for taxon differential abundance analysis, NANOTAXI conducts all-vs-all comparisons for KO terms and MetaCyc pathways, although only one of these comparisons is presented in this study. Gene set enrichment analysis (GSEA) of the ranked differentially abundant KO terms revealed several enriched and depleted pathways in the *E. coli* dominant group compared with the ZymoBIOMICS group. Enriched predicted pathways included those related to stress resistance and persistence, such as beta-lactam resistance and biofilm formation, as well as lipopolysaccharide biosynthesis, a Gram-negative-specific pathway. In contrast, depleted predicted pathways were primarily involved in translation and maintenance of cell wall structure, including ribosome function, aminoacyl-tRNA biosynthesis, peptidoglycan biosynthesis, and teichoic acid biosynthesis, which is unique to Gram-positive bacteria (**Figure 7B**). This pattern indicates that *E. coli* dominance was associated with lower predicted abundance of pathways associated with the Gram-positive members of the ZymoBIOMICS mock community and a reduction in global protein synthesis capacity, while showing higher predicted representation of pathways associated with stress tolerance and cell envelope integrity.

Differential abundance analysis of MetaCyc pathways further supported the GSEA findings. Lipopolysaccharide biosynthesis, which is unique to Gram-negative bacteria, was found to be enriched. In contrast, peptidoglycan biosynthesis, a Gram-positive-specific pathway, was found to be depleted in the differential abundance analysis comparing the *E. coli* dominant group with the ZymoBIOMICS mock community group. Furthermore, lipid A-core biosynthesis, which produces an essential membrane anchor for lipopolysaccharide, was also found to be enriched in the differential abundance analysis (**Figure 7D**).

These results serve as a proof-of-concept, demonstrating that NANOTAXI’s functional module can seamlessly translate taxonomic profiles into predicted metabolic capabilities and perform comparative enrichment analyses, a workflow directly applicable to real microbiome studies where functional potential is of interest.

## 4 Discussion

The 16S rRNA gene serves as a cornerstone phylogenetic marker for profiling microbial communities. While short-read sequencing offers high throughput and accuracy, its limited read length restricts species-level taxonomic resolution and makes it less suited to continuous full-length real-time amplicon profiling. Long-read sequencing platforms, such as those from Oxford Nanopore Technologies (ONT), overcome these constraints by capturing the full-length ∼1.5 kb 16S rRNA gene and generating data in real time. Despite this potential, a significant gap persists: the lack of an integrated, user-friendly platform that connects live Nanopore data streams to comprehensive, production-grade statistical workflows for microbiome analysis.

To address this need, we developed NANOTAXI, an R Shiny application that bridges real-time nanopore sequencing with advanced microbiome analytics. The platform enables automated, parallel processing of barcoded 16S rRNA reads from the earliest stages of a sequencing run, delivering live taxonomic classification, diversity metrics, and interactive visualisations. Crucially, NANOTAXI triggers cohort-level comparative analysis automatically after just five successful classification iterations, allowing group-based statistical comparisons to begin well before the run completes. Upon the user terminating the run, the application finalises data processing. It executes the functional inference module, providing a complete, end-to-end analytical workflow from live monitoring to metabolic prediction within a single, point-and-click interface.

Existing platforms for Nanopore 16S data analysis are often limited to either command-line workflows or predefined pipelines with limited flexibility. For example, EPI2ME’s Nextflow-based 16S pipeline provides real-time taxonomic analysis but offers limited support for downstream statistical analyses commonly used in microbiome research. Furthermore, its command-line interface restricts accessibility for clinicians and researchers with limited bioinformatics experience. NANOTAXI addresses this gap by integrating multiple classification strategies, cohort-level microbiome analyses and functional inference within a unified, interactive environment (Supplementary Table 2).

We validated NANOTAXI using barcoded synthetic communities comprising pooled genomic DNA from clinically relevant bacterial species and the ZymoBIOMICS mock community. The application successfully interfaced with the MinKNOW API to monitor sequencing progress, classify reads in real time and generate progressively updated ecological summaries. The resulting outputs were biologically coherent and consistent with the designed community compositions. Communities dominated by a single species exhibited reduced alpha diversity, while beta-diversity ordination and principal component analysis clearly separated samples according to their expected compositions. Differential abundance analysis correctly identified enrichment of the intended dominant taxa. An unexpected predominance of *E. coli* in a barcode intended for *P. aeruginosa* dominance highlighted NANOTAXI’s utility as a real-time quality-control tool for detecting sample-preparation or pooling errors during sequencing.

Functional inference using PICRUSt2 further demonstrated that taxonomic shifts identified by NANOTAXI translated into biologically meaningful changes in predicted metabolic potential. For example, *E. coli*-enriched samples showed increased representation of lipopolysaccharide biosynthesis pathways and decreased representation of pathways associated with Gram-positive cell wall components, consistent with the underlying taxonomic composition. These results highlight the value of integrating taxonomic profiling with downstream functional prediction, while recognising that PICRUSt2 provides inferred rather than directly measured functional profiles.

To accommodate different analytical objectives and computational constraints, NANOTAXI supports multiple classification tools and reference databases. Assessment of classification performance on the ZymoBIOMICS mock community revealed substantial differences in false positive rates among the four classifiers. Emu showed the highest observed specificity, detecting no false-positive species across all replicates and therefore served as the default high-specificity configuration. MMseqs2 exhibited a low average false-positive abundance of 1.41%, followed by Minimap2 at 3.84%. In contrast, Kraken2 showed the highest false-positive rate, with an average of 16.2% of the abundance attributable to the false-positive species. (Supplementary Figure 5A, Supplementary Table 3). At the genus level, false positive rates decreased across all classifiers: Kraken2 dropped to 4.17%, Minimap2 to 2.42%, and MMseqs2 to 1.29% (Supplementary Figure 5B).

Additionally, we assessed the runtime performance of the classifiers with all available databases to process 500 reads with 4 computational threads, using the default chunk size and computation threads per barcode. Kraken2 was the fastest classifier across all databases, with a mean execution time ranging from 5.65 seconds (GSR) to 9.12 seconds (GTDB) per chunk, making it the most suitable option for continuous near-real-time monitoring under the tested configuration (Supplementary Figure 6A, Supplementary Table 4). Minimap2 showed intermediate performance (11.27–45.14 seconds), followed by MMseqs2 (34.25-115.87 seconds) and Emu (34.42-146.93 seconds). Notably, MMseqs2 consumed substantially more memory (8.4–9.0 GB) than all other classifiers (approximately 1.1 GB), rendering it impractical for resource-limited environments (Supplementary Figure 6B, Supplementary Table 4). Database choice also influenced speed; GSR consistently required longer execution times, likely due to its larger number of entries (90,171), whereas GTDB and REFSEQ (48,752 and 27,352 entries, respectively) were faster across all methods.

These benchmarking results provide practical guidance for selecting an appropriate classification strategy. Kraken2 may be preferred when rapid classification and limited computational resources are the primary considerations, particularly when genus-level resolution is sufficient. For analyses requiring higher species-level specificity, Emu, with its native database, offers the lowest observed false-positive rate, although at the cost of increased computational time. MMseqs2 and Minimap2 offer intermediate alternatives, with MMseqs2 generally requiring substantially more memory. Together, these options allow users to balance speed, taxonomic resolution and hardware requirements according to the objectives of a given study.

The chunked and asynchronous architecture of NANOTAXI ensures that computationally intensive classifiers do not compromise interface responsiveness. Reads are processed in discrete batches, and classification occurs in background processes while the dashboard remains interactive. This design allows users to prioritise higher-resolution classification strategies when sufficient computational resources are available, while maintaining a consistent user experience.

The current version of this study has certain limitations. First, benchmarking was primarily performed using synthetic communities and controlled datasets; further validation across complex clinical and environmental microbiomes will provide a broader assessment of performance under highly diverse conditions. Second, classification speed and memory requirements remain dependent on both hardware specifications and database size, particularly for alignment and mapping-based approaches. Third, although PICRUSt2 extends the platform’s analytical scope, its outputs represent predicted functional potential rather than direct measurements of gene expression or metabolic activity. Finally, as with all reference-based approaches, taxonomic assignments are influenced by the completeness and curation of the underlying databases.

Overall, NANOTAXI represents a significant advance in accessible microbiome bioinformatics. It is a fully automated, one-click solution that demystifies complex analysis pipelines for wet-lab researchers and clinicians. The platform supports two primary modes: real-time analysis during live sequencing and post-run (offline) analysis of previously generated datasets, ensuring broad utility regardless of when the data was acquired. By supporting multiple classification tools and databases, it remains adaptable across clinical, environmental, and forensic applications. Most importantly, NANOTAXI effectively bridges the gap between the real-time data generation capability of nanopore sequencing and the advanced statistical interpretation required for modern microbiome science, empowering a broader community of researchers to leverage long-read amplicon sequencing in their work.

## 5 Conclusions

NANOTAXI provides an integrated GUI-based workflow for real-time and post-run analysis of full-length 16S rRNA nanopore sequencing data. By combining live MinKNOW integration, modular classifier selection, cohort-level microbiome statistics, and functional inference, the platform enables users to move from raw barcoded reads to interpretable biological outputs within a single environment. Validation on synthetic communities demonstrated its ability to recover expected taxonomic patterns, identify sample-level anomalies, and generate coherent functional predictions. NANOTAXI therefore provides an accessible framework for applying long-read amplicon sequencing in clinical, environmental, and experimental microbiome studies.

## 6 Availability and Requirements

NANOTAXI is freely available under the GNU GPL v3 license on GitHub (https://github.com/Nirmal2310/NANOTAXI). The repository includes detailed instructions for setting up the tool and a minimal test dataset.

The bulk fast5 file for the barcoded 16S Nanopore sequencing run is publicly available under the ZENODO platform https://zenodo.org/records/20195479. The step-by-step guide for running the sequencing playback, along with the associated metadata for the run, is available in the NANOTAXI GitHub repository.

## 7 Declarations

### 7.1 Ethics approval and consent to participate

Not Applicable.

### 7.2 Consent for publication

Not Applicable.

### 7.3 Availability of data and materials

Project name: NANOTAXI.

Project Home Page: https://github.com/Nirmal2310/NANOTAXI

Operating System(s): Linux

Programming Language: Bash, R>=4.4.2, Python.

Other Requirements: MinKNOW>=24.06.10, and Conda >=4.12.0.

License: GNU GPL v3.0.

## Author Contributions

IG conceived the concept and supervised the findings. NSM created the application, performed the analysis and drafted the manuscript. KC prepared the sequencing library and ran the sequencing. All authors wrote, read and approved the final manuscript.

