## Supplementary Figures for "NANOTAXI: A Shiny-Based GUI for Real-Time Classification and Analysis of 16S rRNA Nanopore Reads"

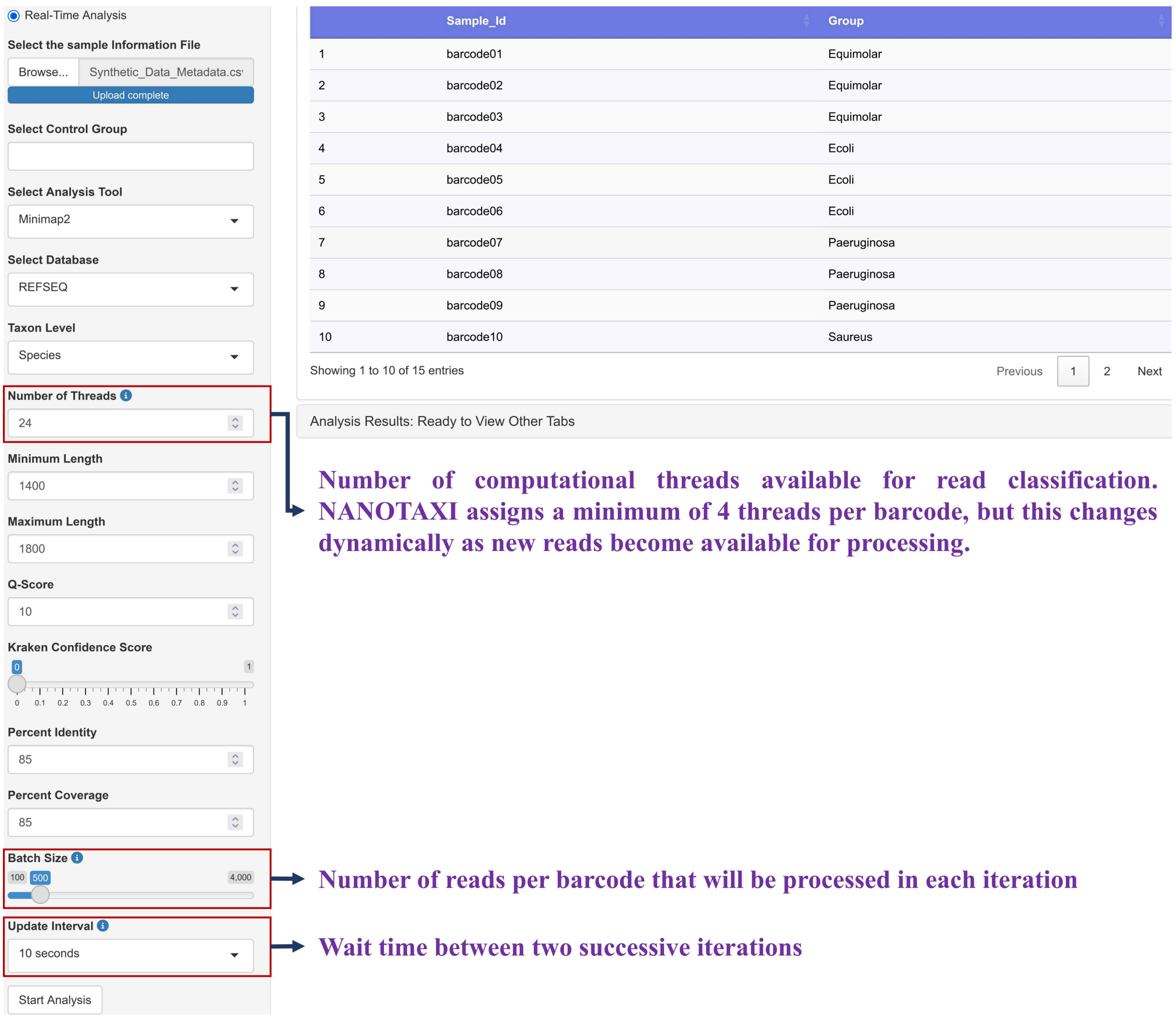


**Supplementary Figure 1: A general overview of the options provided by NANOTAXI for real-time analysis.** Detailed information on some key parameters is provided in the figure.

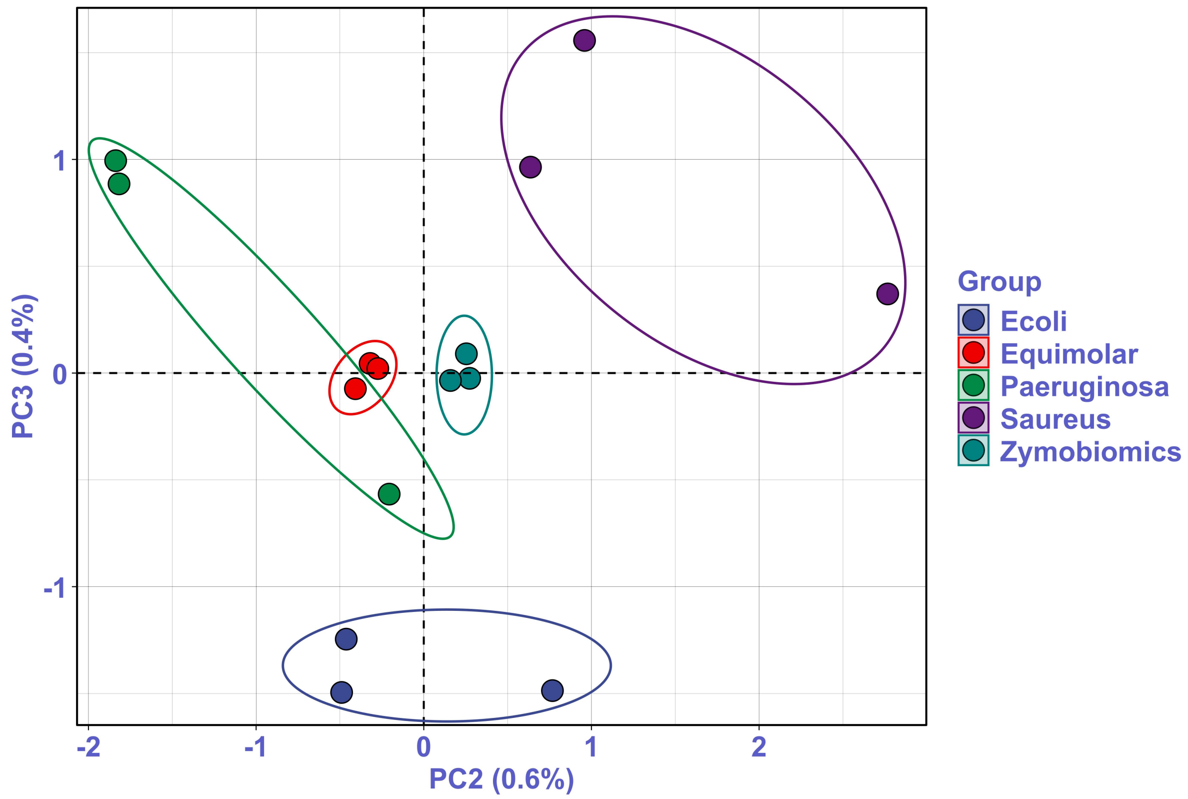


**Supplementary Figure 2: Principal Coordinate Analysis (PCoA) ordination plot of the second and third principal coordinates.** The percentage variance explained is displayed on each axis.

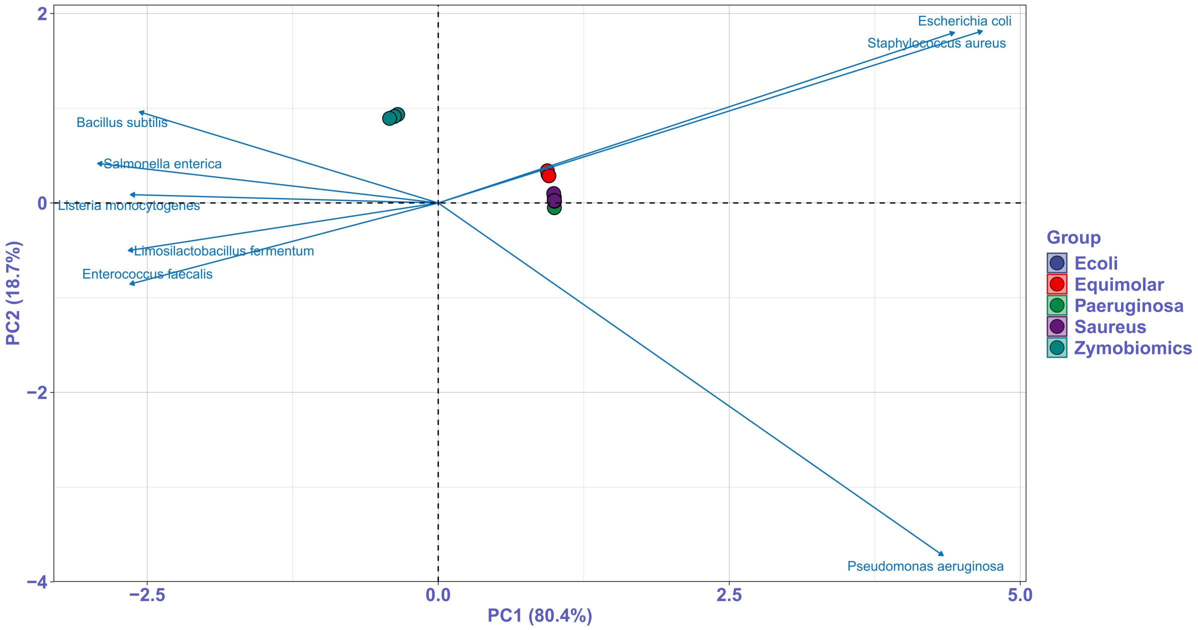


**Supplementary Figure 3: Principal Component Analysis (PCA) ordination of CLR-transformed data.** The top taxa (default: 5; adjustable to 25) overlaid as vectors to indicate taxa driving group separation along the first two principal components.

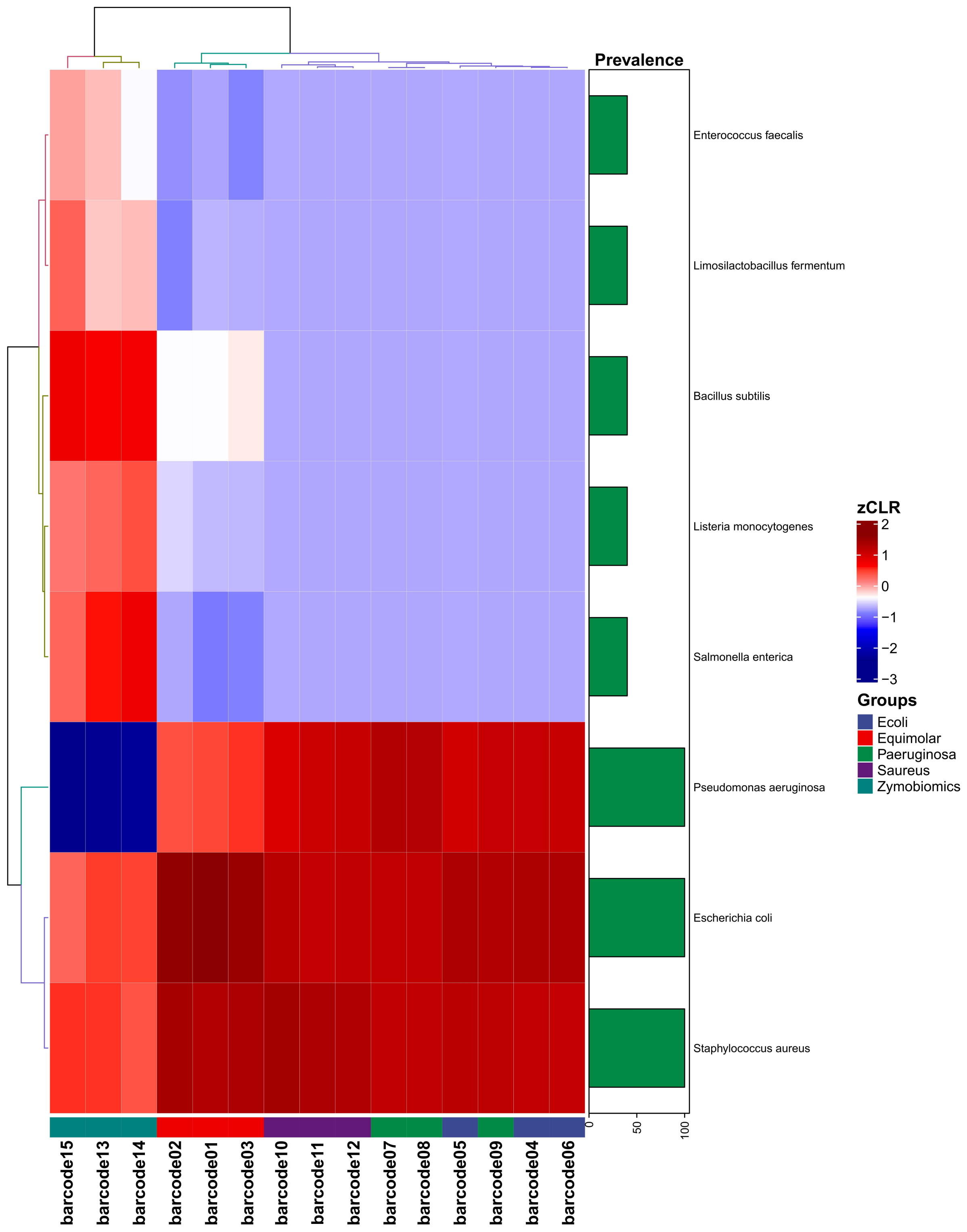


**Supplementary Figure 4: Heatmap of z-score standardised CLR-transformed abundances.** Rows (taxa) and columns (barcodes) are hierarchically clustered using Euclidean distance with complete linkage. Barcodes are annotated by experimental group, and a sidebar displays taxon prevalence.

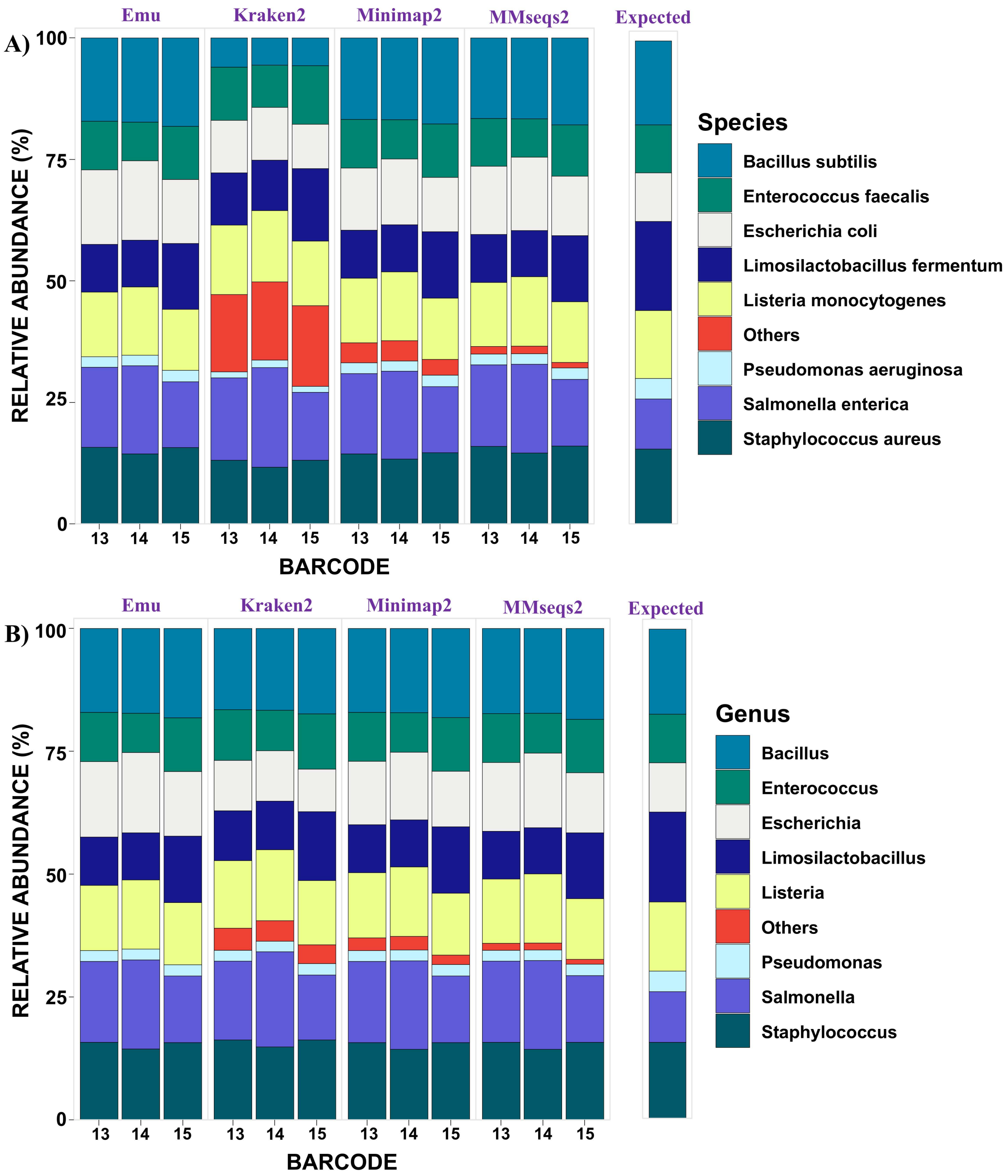


**Supplementary Figure 5: Comparison of classification accuracy of all four classification methods over the Zymobiomics mock community.** A) Stacked bar plot showing the relative abundance of the classified species based on the four methods. B) Stacked bar plot showing the relative abundance of the classified genus based on the four methods.

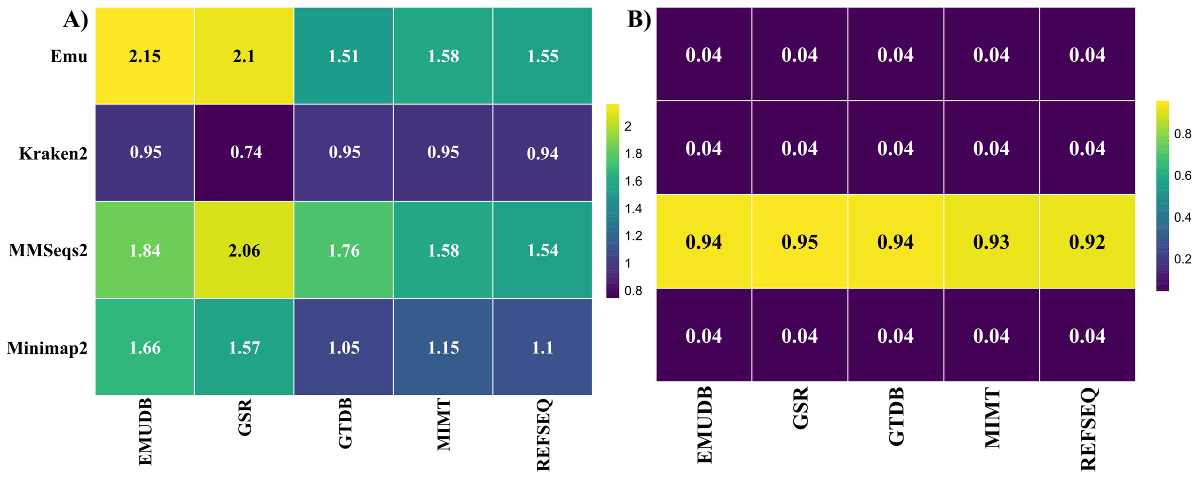


**Supplementary Figure 6: Comparison of execution time and peak memory usage for each classification tool and database combination.** A) Heatmap showing the median execution time (seconds) by each combination of classification tool and database to process a single chunk (500 reads) in $\log_{10}$ scale. B) Heatmap showing the median peak memory usage (GB) by each combination of classification tool and database to process a single chunk in $\log_{10}$ scale.
